# CART neuropeptide modulates the extended amygdalar CeA-vBNST circuit to gate expression of innate fear

**DOI:** 10.1101/096610

**Authors:** Abhishek Rale, Ninad Shendye, Devika S Bodas, Nishikant Subhedar, Aurnab Ghose

## Abstract

Innate fear is critical for the survival of animals and is under tight homeostatic control. Deregulation of innate fear processing is thought to underlie pathological phenotypes including, phobias and panic disorders. Although central processing of conditioned fear has been extensively studied, the circuitry and regulatory mechanisms subserving innate fear remain relatively poorly defined.

In this study, we identify cocaine- and amphetamine-regulated transcript (CART) neuropeptide signaling in the central amygdala (CeA) - ventral bed nucleus of stria terminalis (vBNST) axis as a key modulator of innate fear expression. 2,4,5-trimethyl-3-thiazoline (TMT), a component of fox faeces, induces a freezing response whose intensity is regulated by the extent of CART-signaling in the CeA neurons. Abrogation of CART activity in the CeA attenuates the freezing response and reduces activation of vBNST neurons. Conversely, ectopically elevated CART signaling in the CeA potentiates the fear response concomitant with enhanced vBNST activation. We show that local levels of CART signaling modulate the activation of CeA neurons by NMDA receptor-mediated glutamatergic inputs, in turn, regulating activity in the vBNST.

This study identifies the extended amygdalar CeA-vBNST circuit as a CART modulated axis encoding innate fear. CART signaling regulates the glutamatergic excitatory drive in the CeA-vBNST circuit, in turn, gating the expression of the freezing response to TMT.

## 1. Introduction

Exposure to threatening stimuli evokes a constellation of responses aimed at self-preservation. Genetically ingrained mechanisms engender spontaneous fear, independent of earlier experience, and offer a unique opportunity to dissect an emotion – from its arousal to behavioral end point. While the onset, intensity, persistence and extinction kinetics of these innate responses are tightly regulated, dysregulation of the underlying system may lead to neurological conditions like post-traumatic stress disorders, phobias and panic disorders. Unimodal predator cues, like 2,4,5-trimethyl-3-thiazoline (TMT; an ethologically relevant fear-inducing odorant derived from fox faeces) have been used to delineate the neuroanatomical underpinnings of innate fear (Day et al., 2004; Rosen et al., 2015; Silva et al., 2016; Takahashi, 2014).

TMT is sensed by discrete neurons of the nasal epithelium and Gruenenberg ganglia, which project to the main olfactory bulb as well as the accessory olfactory bulb (Brechbuhl et al., 2013; Kobayakawa et al., 2007; Matsumoto et al., 2010). Downstream to the medial or accessory olfactory bulbs, the TMT generated information is known to transit via the cortical nucleus of the amygdala (CoA) (Root et al., 2014) and medial nucleus of the amygdala (MeA) (Muller and Fendt, 2006). How the information transits from the CoA/MeA to motor output areas like the periaqueductal grey (PAG) remains unclear. A possible route may involve central nucleus of the amygdala (CeA) – ventral bed nucleus of the stria terminalis (vBNST) connectivity that, in turn, communicates to the PAG possibly via specific hypothalamic nodes (Motta et al., 2009; Pagani and Rosen, 2009). A tight coordination between the CeA and BNST is emerging as a major regulatory node in the processing of fear and anxiety in rodents and primates (Fox et al., 2015; Shackman and Fox, 2016). Previous studies based on rodents as well as primates implicate CeA in a variety of fear responses triggered by predator or predator cues (Day et al., 2004; Kalin et al., 2004). Suppression of activity of serotonin-2A receptor expressing neurons of the CeA has been shown to mediate innate fear induced by an artificial TMT-derivative (Isosaka et al., 2015). Exposure to ferret resulted in an increase in the secretion of CRF in the CeA of rat (Merali et al., 2001). Studies from our laboratory and others implicate neuronal activation in the CeA in response to TMT (Butler et al., 2011; Sharma et al., 2014), while silencing of the vBNST by muscimol abolished TMT-induced freezing (Fendt et al., 2003).

In contrast to conditioned fear, our understanding of the modulatory control of innate fear is limited. The amygdalar circuitry has emerged as a central regulatory hub for conditioned fear. A range of agents, inclusive of fast-acting neurotransmitters like GABA, glutamate, dopamine and serotonin and neuropeptides like corticotropin-releasing factor (CRH), opioid peptides, neuropeptide Y, thyrotropin-releasing hormone, calcitonin gene-related peptide and vasopressin influence fear conditioning (Davis and Whalen, 2001; Schulkin et al., 2005; Shionoya et al., 2013; Spannuth et al., 2011; Tasan et al., 2016). However, little is known about the modulatory processes associated with innate fear. CRF, somatostatin and opioids have been implicated in innate fear processing (Asok et al., 2013; Asok et al., 2016; Figueiredo et al., 2003; Nanda et al., 2008; Roseboom et al., 2007; Wilson and Junor, 2008), and the underlying modes of action and neuroanatomical substrates are just beginning to be understood.

Studies from our laboratory have implicated the neuropeptide CART as an important player in the processing of innate fear within the CeA. Exposure to cat or TMT induced robust freezing in rats, which was dependent on CART signaling (Sharma et al., 2014; Upadhya et al., 2013). In this study, we uncover a CART signaling-sensitive extended amygdalar CeA-vBNST circuitry in TMT-induced fear processing. CART potentiates NMDA-R-dependent excitatory drive in the CeA-vBNST axis and exerts regulatory control on TMT-induced freezing.

## 2. Materials and methods

### 2.1. Animals

Adult male Sprague-Dawley rats weighing 200–220 g at the time of surgery were used. All the rats were maintained on a 12 hr light / dark cycle, at controlled room temperature of 25± 2°C with food and water available *ad libitum*. The bedding of the cages was changed every week. In order to obviate novelty related stress, all rats were habituated for five days to handling, laboratory conditions and to the test chamber. All experimental protocols were approved by the Institutional Animal Ethical Committee (IAEC) constituted by the CPCSEA, Govt. of India.

### 2.2. Surgery

Stereotaxic surgery and implantation of cannula were carried out according to previously described protocols (Sharma et al., 2014). Briefly, the rats were anaesthetized with intraperitoneal (i.p.) ketamine (60 mg/kg, Aqua Fine Injecta, India) and xylazine (10 mg/kg, Stanex, India) injection. Hair depilator (Anne French, Wyeth, India) was applied to the head to remove hair. Each rat was mounted on the stereotaxic frame with blunt ear bars (Stoelting, USA) and a mid-sagittal incision was made in the scalp to expose the skull. Two stainless steel guide cannulae were implanted bilaterally targeted at the CeA using the stereotaxic coordinates -1.9 mm caudal, ± 4.0 mm lateral and -7.8 mm ventral to the bregma and secured to the skull with anchoring screws and dental cement (DPI-RR cold cure, acrylic powder, Dental Products of India, India). After surgery and between testing, dummy cannulas were inserted into the guide cannulas to prevent occlusion. The animals were placed in separate cages to avoid damage to the guide and dummy cannulae. All rats were allowed one week to recover prior to the start of behavioral testing. Only the rats showing quick recovery and no signs of infection were included in the study. The animals were divided randomly into different groups (n = 6 in each) and habituated to the testing environment for five days.

Post-necropsy, the brain sections were examined for proper placement of the cannulae (Supplementary Fig S1). The data drawn from the animals with both the cannulae in the target region were considered for analysis.

### 2.3. Microinjections

For microinjections, the injection cannulae (fabricated in-house; internal diameter 0.16 mm, outer diameter 0.31 mm) connected via PE-10 polyethylene tubing to a microliter syringe (10 μl, Hamilton, USA) and extending 0.5 mm beyond the guide cannulae (fabricated in house as described earlier (Kokare et al., 2011); internal diameter 0.36 mm, outer diameter 0.5 mm) targeting the CeA were used and rats were bilaterally administered different agents according to their treatment group. The control group was bilaterally injected with 0.5 μl of artificial cerebrospinal fluid (aCSF; 140 mM NaCl, 3.35 mM KCl, 1.26 mM CaCl_2_, 1.15 mM MgCl_2_, 0.3 mM NaH_2_PO_4_, 1.2 mM Na_2_HPO_4_ (pH 7.4) over a period of 5 min. Similarly, other groups received different treatments such as non-immune serum (NIS; 0.1% bovine serum albumin in aCSF; 0.5 μl/side); CART antibody (1:500 in NIS; 0.5 μl/side; gift from Drs. Lars Thim and Jes Clausen, Novo Nordisk, Denmark); CART peptide (10 ng in 0.25 μl/side; gift from Drs. Lars Thim and Jes Clausen, Novo Nordisk, Denmark); lidocaine hydrochloride (2% solution; AstraZeneca) and MK 801, a non-competitive NMDAR antagonist (5 μg in 0.5 μl/side, diluted in aCSF; Tocris) bilaterally in the CeA. For administration of CART antibody, CART peptide or matching controls in the CeA, the animals were injected 15 mins prior to behavioral testing. MK801 and control aCSF were injected 5 mins prior to behavior testing, while lidocaine and buffered saline was injected 10 mins prior. For double injections, aCSF or MK801 was bilaterally injected into the CeA followed by a second bilateral injection of CART peptide after 5 mins. Behavioral tests were conducted 15 mins after the second injection.

An additional experiment was conducted to investigate the effect of CART treatment per se on the CeA neurons. Sodium thiopental (60 mg/ml; i.p.) anaesthetized rats were stereotactically injected with CART peptide (10 ng dissolved in 0.5 μl aCSF) bilaterally in the CeA using a 31-gauge needle. Following an interval of 30 mins, the animals were perfused transcardially and subjected to immunofluorescence analysis (see below).

### 2.4. Exposure of rat to TMT and behavior assessment

Behavioral tests were carried out as described earlier (Sharma et al., 2014). In this publication, we characterized the specificity of TMT to induce fear and neuronal activation in the CeA and the vBNST as opposed to butyric acid - a non-specific, aversive response to a noxious odorant. Briefly, rats were habituated to the Plexiglas test chamber having dimensions 8.6 × 8.6 × 20 cm (Wallace and Rosen, 2001) following recovery and equipped with two doors at opposite ends (8.6 × 8.6 cm) each having a 6 × 6 cm opening covered by the filter paper. The animals were habituated for 10 mins each day for 5 days. On the 6^th^ day, fifteen minutes after the injections, two filter papers, each coated with 35 μl of TMT were taped over the two openings and the rat was introduced into the test chamber. The behavior of the rat was monitored for a period of 20 min. During the test period, the freezing behavior (absence of all movements except those required for respiration) was recorded and analysed using Noldus Ethovision video tracking system (Netherlands). During the test period, the videos were acquired at 8.33 frames per second. The analysis was conducted by averaging 20 frames. 5% change in body area was set as the threshold to quantify freezing behavior. The data acquisition was by a skilled individual blind to the treatments. The data on freezing are represented as percent of total recorded time.

### 2.5. Immunohistochemistry

The protocol described in our earlier study was employed (Sharma et al., 2014). Thirty minutes after TMT exposure, the rats were anesthetized (sodium thiopental; 60 mg/kg; i.p.) and perfused transcardially using saline followed by chilled 4% paraformaldehyde (PFA) in 0.1 M phosphate buffer (pH 7.4). The brains were post-fixed in 4% PFA overnight and transferred to 30% sucrose solution at 4°C for cryoprotection. The brains were serially sectioned on a cryostat in a coronal plane at the 30 μm thickness and stored in 50% glycerol in PBS at 4°C. Free-floating sections were rinsed in PBS, incubated in the blocking solution containing polyclonal anti-Fos antibody (1:1000, Santa Cruz, USA) for 24 h in a humid atmosphere at 4°C. The sections were then washed and incubated with anti-rabbit Alexa Fluor 488 secondary antibody (A-11008; Invitrogen, Carlsbad, CA) for 2 hr. Further, sections were mounted in a glycerol-based mounting medium (70% glycerol, 0.5% N-propyl-gallate, 20 mM Tris, pH 8.0) containing 4,6-diamidino-2-phenylindole (DAPI; 0.01 mg/ml) and observed under an epifluorescence microscope (AxioImager Z1, Carl Zeiss, Germany) or imaged using a laser scanning confocal microscope (Zeiss LSM 710, Carl Zeiss, Germany). ImageJ was used to adjust the size, contrast, and brightness of the micrographs. Inkscape (ver. 0.91) was used to prepare the panels and diagrammatic representations. In order to ensure reliable comparisons across different groups and maintain stringency in tissue preparation and staining conditions, all brain sections were processed concurrently under identical conditions.

### 2.6. Morphometric analysis

Morphometric analysis was carried out according to the protocol described in our earlier study (Sharma et al., 2014). Briefly, the number of Fos-expressing cells were counted from eight sections containing both the sides of vBNST and dlBNST regions (A.P -0.12 mm to -0.36 mm with reference to bregma), drawn from each of the six brains in each group (Supplementary Fig S1). The morphometric scoring was by a skilled individual blind to the treatments. The cell numbers were subjected to Abercrombie’s correction to avoid overestimation using the equation *N = (p × T)/(T + d)*, where *N* is the corrected cell number, *T* is the thickness of section, *p* is the actual profile count and *d* is the mean nuclear diameter.

### 2.7. Statistical Analysis

Behavior and morphometric data analyses were performed using Mann-Whitney test. All values are expressed as mean ± SEM of the group and differences were considered significant at *p*<0.05. Graphs were plotted using the GraphPad Prism 5.0 statistical software.

## 3. Results

### 3.1. CART signaling in the CeA-vBNST circuit modulates expression of TMT-induced innate fear

As the extended amygdalar CeA-BNST circuit has been implicated in processing innate fear (Schulkin et al., 2005), we tested if CART signaling modulated the activity of this circuit following exposure to TMT. To directly test if CeA activity regulates innate fear and vBNST activation, we silenced CeA neurons by stereotaxically administering lidocaine in the CeA. Compared to controls, lidocaine injected animals showed reduced freezing in response to TMT (Fig 1A; p = 0.0022; n = 6). Our previous work had implicated activation of vBNST neurons by TMT (Sharma et al., 2014). Fos expression is a common surrogate readout of recent neuronal activation. We evaluated the activity of vBNST neurons, upon silencing of the CeA, employing induction of Fos. Lidocaine administration to the CeA was found to attenuate TMT-induced Fos activation in the vBNST neurons (Fig 1B-F; p = 0.0022; n = 6).

**Figure 1.**
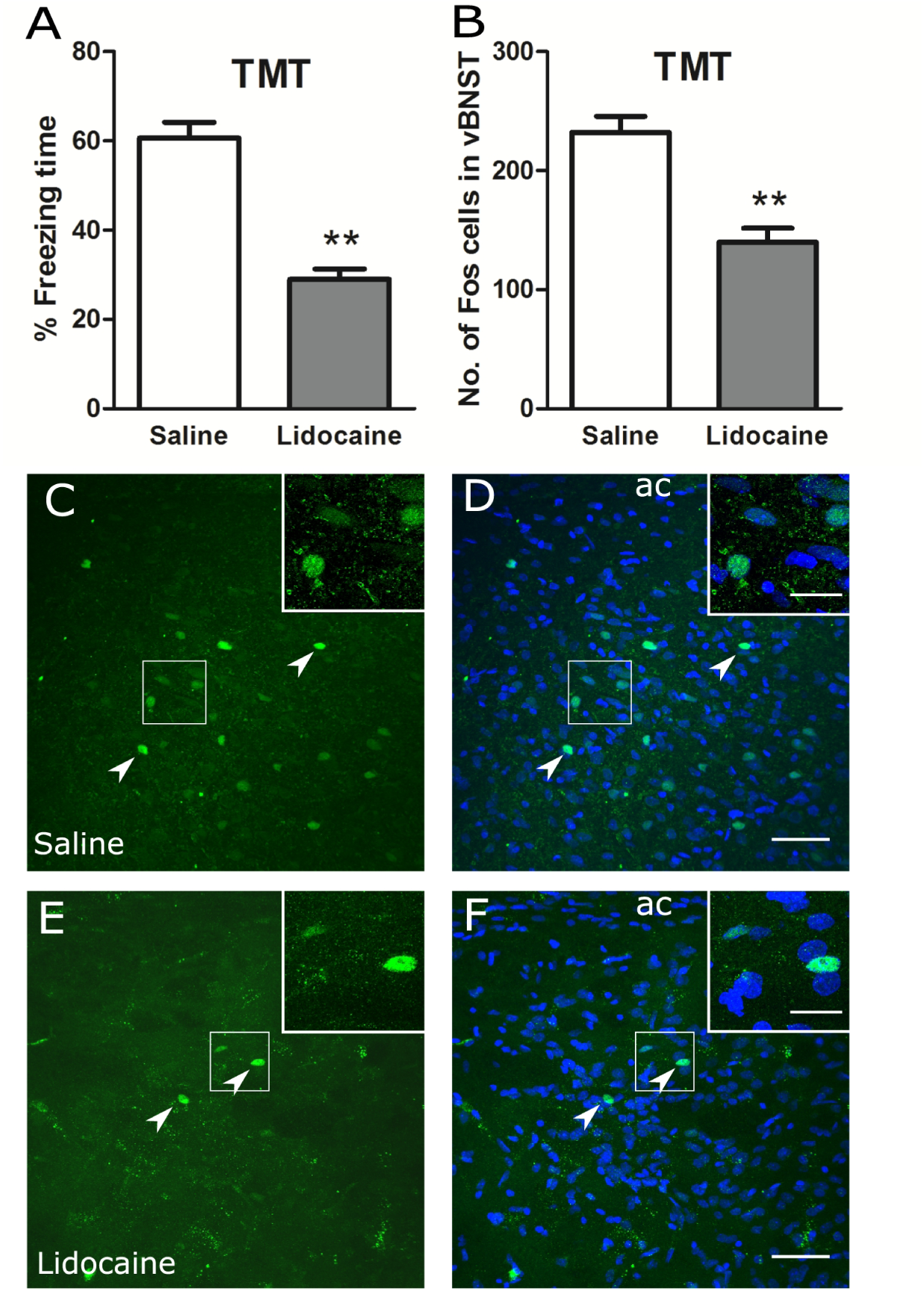
CeA mediates TMT-induced freezing and activation of the vBNST neurons. The effects of intra-CeA administration of buffered saline or lidocaine on percent time spent freezing (A) and the number of Fos-positive cells in the vBNST (B) following TMT exposure are represented as means ± SEM. Representative micrographs of the vBNST region from saline (C-D) and Lidocaine (E-F) injected animals showing Fos staining (C,E) and overlay (D,F) of Fos (green; arrowheads) and DAPI (blue). ac, anterior commissure. The data were analyzed by Mann-Whitney test. *N* = 6 animals in each group. ** *p* < 0.01. Scale bar: 50 μm, (C) - (F); 20 μm, insets.

To test the contribution of CART signaling in processing TMT-induced innate fear responses at the CeA, immunoneutralization of endogenous CART activity was employed. Stereotactic delivery of neutralizing antibody against CART neuropeptide in the CeA of TMT-exposed rat attenuated the freezing response (Fig 2A; p = 0.0022; n = 6) compared to animals injected with the non-immune serum (NIS). This is along the lines of our previous report implicating CART activity in the CeA in processing TMT-induced fear responses (Sharma et al., 2014). We next tested if attenuation of CART signaling in the CeA impacted neural activation in the vBNST. Strikingly, rats injected with CART antibody in the CeA showed reduced Fos induction in the vBNST in response to TMT compared to NIS injected animals (Fig 2B-F; p = 0.0043; n = 6).

**Figure 2.**
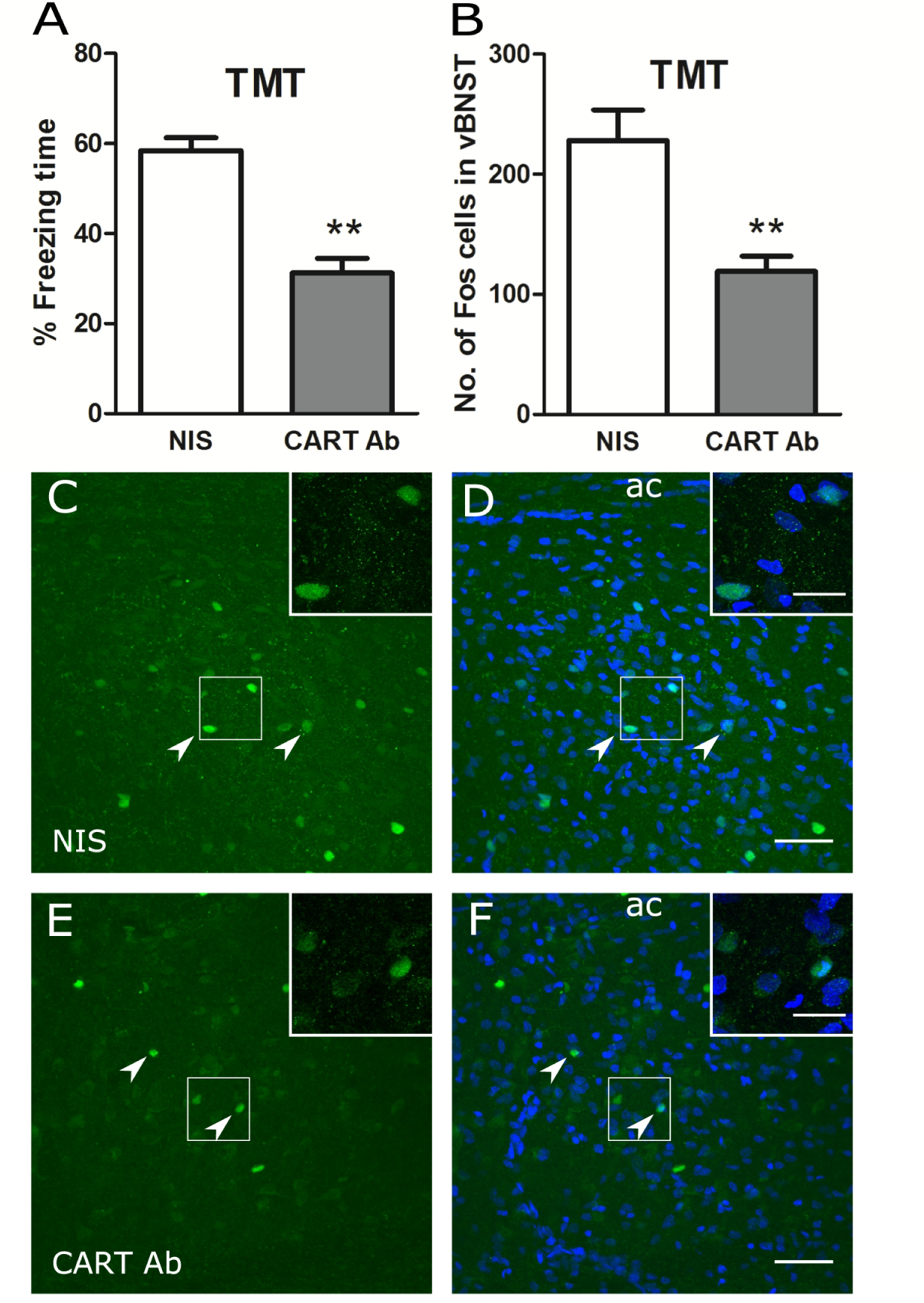
CART signaling in the CeA regulates TMT-induced freezing and activation of the vBNST neurons. The effects of intra-CeA administration of non-immune serum (NIS) or CART antibody (CART Ab) on percent time spent freezing (A) and the number of Fos-positive cells in the vBNST (B) following TMT exposure are represented as means ± SEM. Representative micrographs of the vBNST region from NIS (C-D) and CART Ab (E-F) treated animals showing Fos staining (C,E) and overlay (D,F) of Fos (green; arrowheads) and DAPI (blue). ac, anterior commissure. The data were analyzed by Mann-Whitney test. *N* = 6 animals in each group. ** *p* < 0.01. Scale bar: 50 μm, (C) - (F); 20 μm, insets.

These results, in line with our previous reports (Sharma et al., 2014; Upadhya et al., 2013), confirm the role of CART in processing innate fear in the CeA. Further, we identify a circuitry involving CeA mediated activation of vBNST under the modulatory influence of CART neuropeptide in TMT-induced fear processing.

### 3.2. Exogenous CART peptide in the CeA intensifies TMT-induced fear responses and promotes vBNST activation

To test the role of CART signaling in the CeA-vBNST circuit, CART peptide was administered stereotaxically to the CeA. Compared to aCSF treated animals, CART infused rats showed a significantly prolonged freezing response (Fig 3A; p = 0.0087; n = 6). Fos induction in the vBNST was increased in rats treated with exogenous CART peptide compared to those receiving aCSF in the CeA (Fig 3B-F; p = 0.0043; n = 6). These results highlight the importance of CART signaling in fear processing in the CeA-vBNST axis.

**Figure 3.**
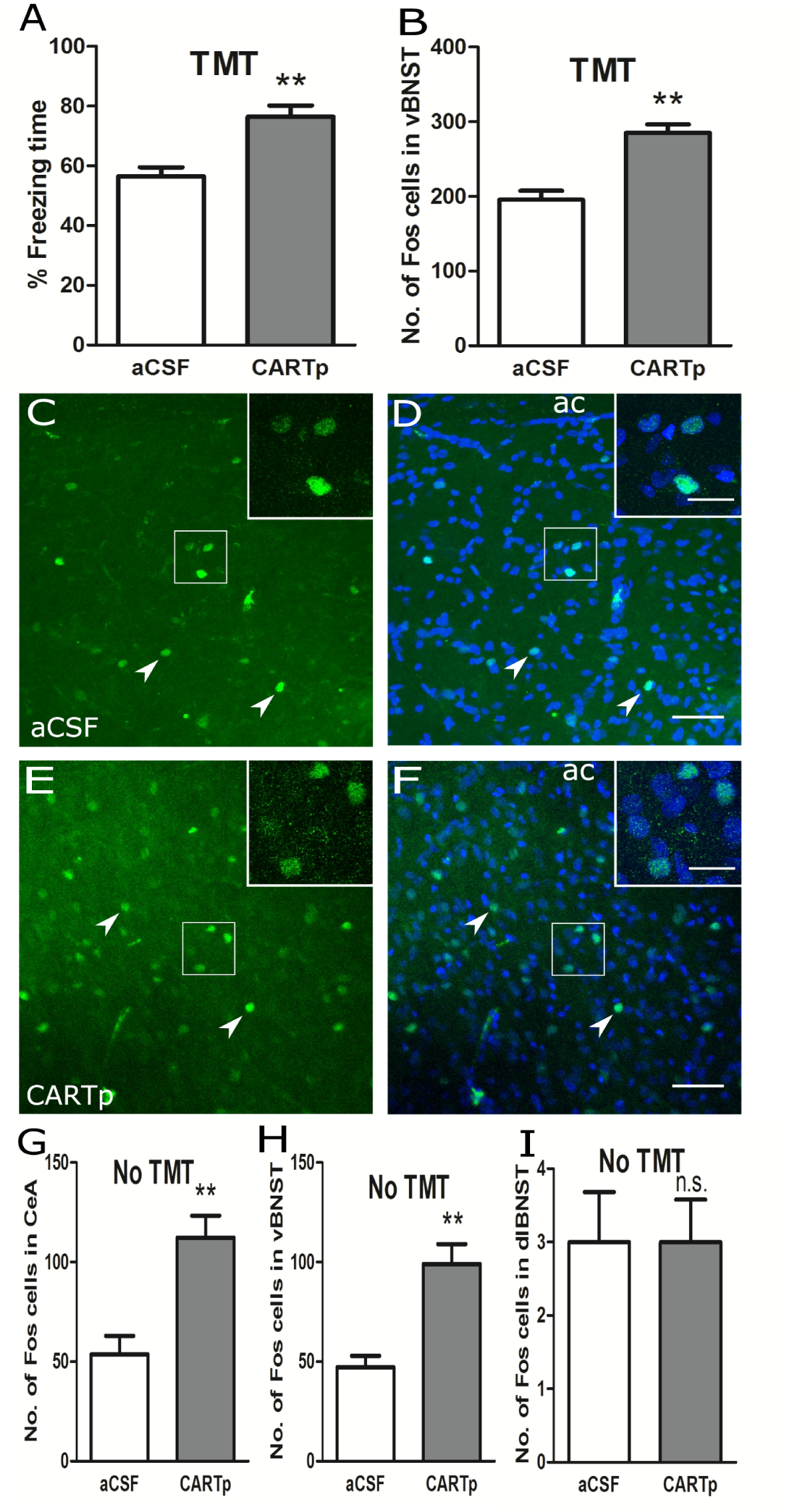
CART peptide intensifies TMT-induced freezing and activation of the vBNST and CeA neurons. The effects of intra-CeA administration of artificial cerebrospinal fluid (aCSF) or CART peptide (CARTp) on percent time spent freezing (A) and the number of Fos-positive cells in the vBNST (B) following TMT exposure are represented as means ± SEM. aCSF or CARTp were administered 15 min prior to TMT exposure. Representative micrographs of the vBNST region from aCSF (C-D) and CARTp (E-F) injected animals showing Fos staining (C, E) and overlay (D, F) of Fos (green; arrowheads) and DAPI (blue). ac, anterior commissure. Intra-CeA administration of CARTp in anesthesized rats, not exposed to TMT, on the number Fos-positive cells in the CeA (G), vBNST (H) and dlBNST (I). The data were analyzed by Mann-Whitney test. *N* = 6 animals in each group (A-B) and *N* = 6 amygdalae in each group (G-I). ** *p* < 0.01. Scale bar: 50 μm, (C) - (F); 20 μm, insets.

We also tested if exogenously applied CART peptide in the CeA could directly alter the excitability of the CeA-vBNST axis in the absences of TMT exposure. Number of Fos-positive neurons increased in both the CeA (Fig 3G; p=0.0043; n=6) and the vBNST (Fig 3H; p = 0.0050; n = 6) when CART peptide was introduced even in the absence of TMT. However, no change in Fos immunoreactive cell population was observed in the dlBNST (Fig 3I; n = 6).

The data drawn from CART immunoneutralization, administration of CART peptide (with or without TMT) and silencing of CeA neurons suggest a strong correlation between CART activity in the CeA, activation of vBNST and expression of TMT-induced fear. The strength of CART signaling in the CeA appears to regulate the intensity of vBNST activation, in turn, gating the expression of innate fear. The intensification of TMT-induced freezing in response to exogenous CART peptide is in line with previous experiments using a live cat as the fear-inducing cue (Upadhya et al., 2013).

### 3.3. NMDA-R activity in the CeA mediates TMT-induced fear processing

With a view to test the involvement of glutamatergic signaling in the CeA in TMT-induced fear, we blocked the NMDA-R activity with MK801, a non-competitive antagonist of NMDA-R. Administration of MK801 directly into the CeA, attenuated the TMT-induced the freezing response (Fig 4A; p = 0.0043; n = 6). MK801 treated animals also showed reduced Fos expression in the vBNST, compared to aCSF controls (Fig 4B-F; p = 0.0087; n = 6). The results underscore the role of glutamatergic inputs, acting via NMDA-R, in conveying the fear information over the CeA-vBNST circuit.

**Figure 4.**
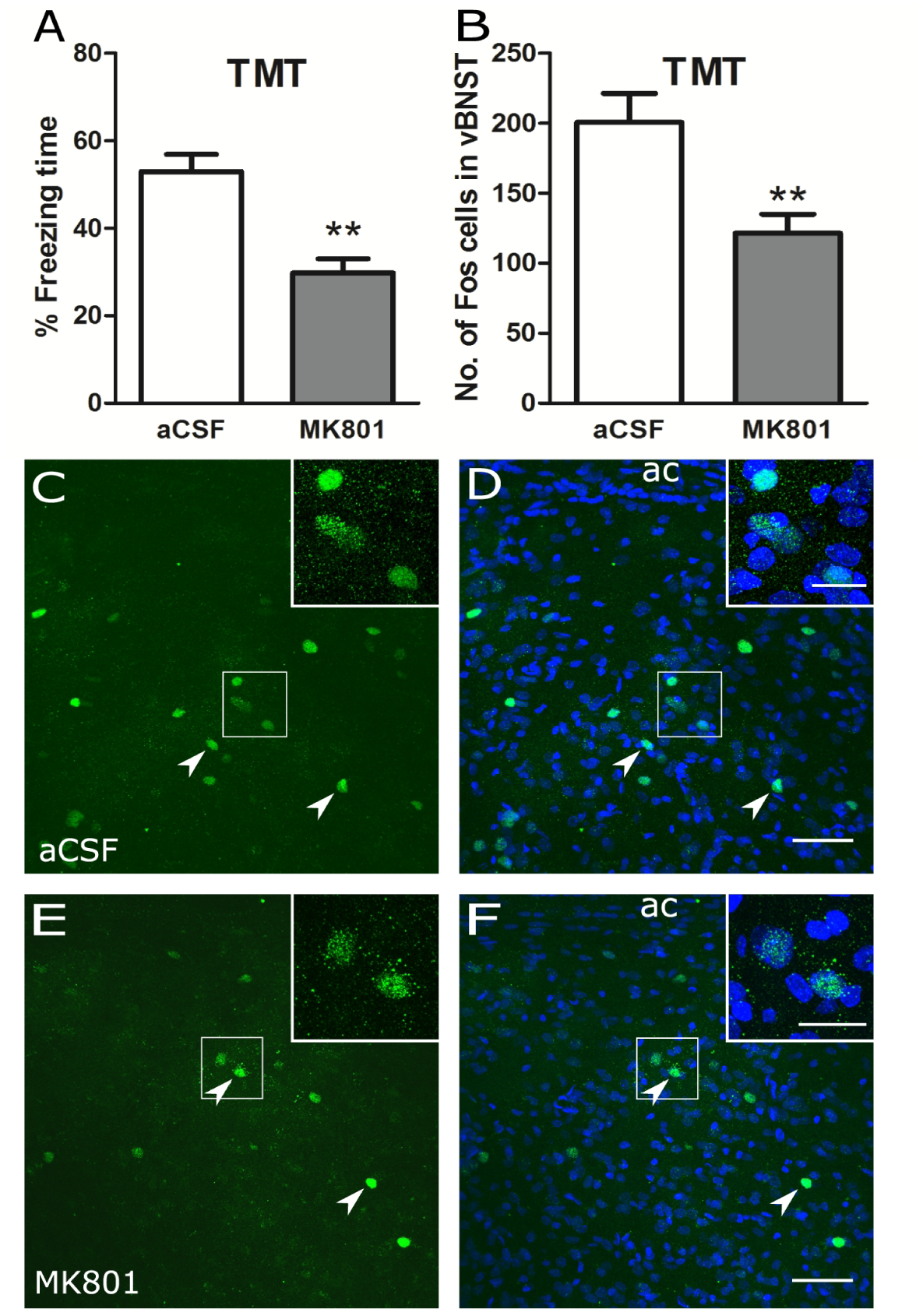
NMDA-R signaling in the CeA mediates TMT-induced freezing and activation of the vBNST neurons. The effects of intra-CeA administration of artificial cerebrospinal fluid (aCSF) or MK801 on percent time spent freezing (A) and the number of Fos-positive cells in the vBNST (B) following TMT exposure are represented as means ± SEM. The agents were administered 5 mins prior to TMT exposure. Representative micrographs of the vBNST region from aCSF (C-D) and MK801 (E-F) treated animals showing Fos staining (C, E) and overlay (D, F) of Fos (green; arrowheads) and DAPI (blue). ac, anterior commissure. The data were analyzed by Mann-Whitney test. *N* = 6 in each group. ** *p* < 0.01. Scale bar: 50 μm, (C) - (F); 20 μm, insets.

### 3.4. CART function in the CeA is mediated by NMDA-R signaling

Based on the above results, we next tested if fear intensification by exogenous CART peptide is mediated by NMDA-R signaling. Animals pretreated with MK801 in the CeA were evaluated for fear potentiation by exogenous CART. CART peptide induced increase in freezing, in response to TMT, was attenuated in MK801 pretreated animal (Fig 5A; p = 0.0043; n = 6). However, vehicle control failed to attenuate the CART peptide augmented response to TMT. Increased expression of Fos in the vBNST, following CART peptide infusion and TMT exposure, was also attenuated upon blocking NMDA-R by MK801 (Fig 5B-F; p = 0.0022; n = 6).

**Figure 5.**
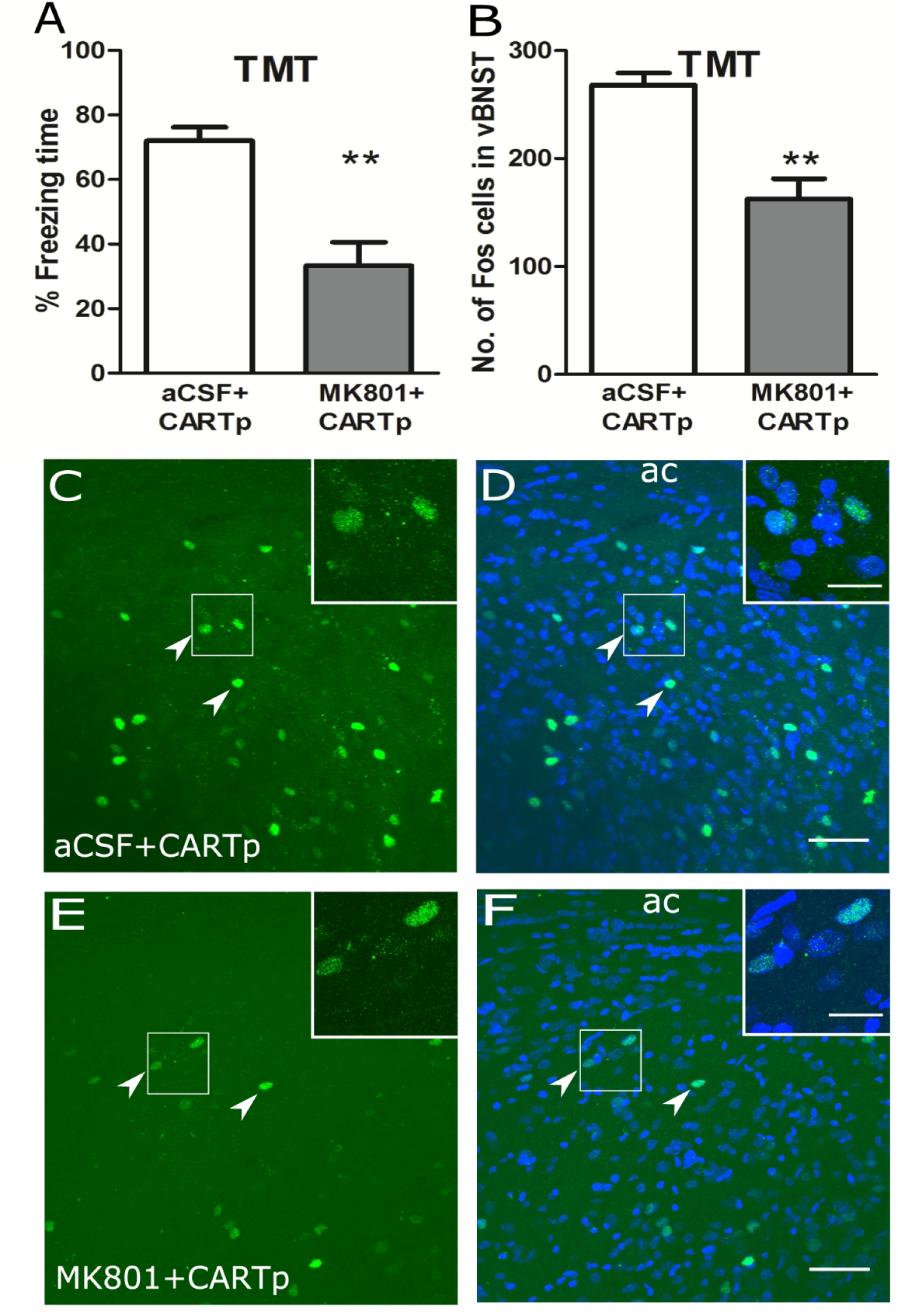
CART activity in the CeA mediates TMT-induced freezing and activation of the vBNST neurons via NMDA-R signaling. The effects of intra-CeA administration of artificial cerebrospinal fluid (aCSF) or MK801 followed by CARTp on percent time spent freezing (A) and the number of Fos-positive cells in the vBNST (B) following TMT exposure are represented as means ± SEM. MK801 or aCSF was administered 15 mins prior to CARTp, followed by TMT exposure after another 5 mins. Representative micrographs of the vBNST region from aCSF + CARTp (C-D) and MK801 + CARTp (E-F) injected animals showing Fos staining (C, E) and overlay (D, F) of Fos (green; arrowheads) and DAPI (blue). ac, anterior commissure. The data were analyzed by Mann-Whitney test. *N* = 6 in each group. ** *p* < 0.01. Scale bar: 50 μm, (C) - (F); 20 μm, insets.

Taken together, these results suggest that CART activity increases the excitatory drive from CeA to vBNST via potentiation of the NMDA-R activity. Modulation of vBNST activation by CART signaling in the CeA regulated the intensity of TMT-induced freezing behavior.

## 4. Discussion

TMT has been widely used to investigate the processing of innate and unconditioned fear. It serves as a reliable, unimodal and ethologically relevant odorant cue that induces fear in rodents. Fear can be quantified in terms of time the animal shows freezing in response to TMT. In this study, we use TMT to characterize the modulation of innate fear by CART neuropeptide.

### 4.1. CART mediated modulation of the CeA-vBNST axis is central to innate fear

The role of specific amygdalar subregions is well characterized in conditioned fear but not for innate fear. Excitotoxic lesions of the basal (BA) and lateral (LA) nuclei of the amygdala did not affect TMT-induced freezing (Wallace and Rosen, 2001). However, muscimol based inactivation revealed the involvement of the basolateral (BLA) and medial amygdala (MeA) (Muller and Fendt, 2006), but not of the LA (Fendt et al., 2003). The amygdalar cortex (CoA) has also been implicated in fear response to TMT (Root et al., 2014). Further downstream, the role of CeA in TMT- and cat-induced fear has been demonstrated in the context of CART neuropeptide signaling (Sharma et al., 2014; Upadhya et al., 2013).

Elevated expression of Fos protein, c-fos mRNA and early growth response 1 (egr-1) mRNA in response to TMT was reported in the rat CeA (Asok et al., 2013; Sharma et al., 2014). These studies also showed increased CART and CRH expression in the CeA underscoring a functional role of CeA in TMT-induced fear processing. On similar lines, ferret odor increased CRH and c-fos expression in the rat CeA (Butler et al., 2011; Merali et al., 2001).

The BNST is a major downstream target of the CeA neurons (Sah et al., 2003; Shackman and Fox, 2016). Several studies have implicated the BNST as a central node in processing innate fear. The neurons of the BNST showed Fos induction on exposure of the rat to predator cues, including TMT (Asok et al., 2013; Campeau et al., 2008; Day et al., 2004; Dielenberg et al., 2001; Figueiredo et al., 2003; Janitzky et al., 2009; Masini et al., 2005; McGregor et al., 2004; Sharma et al., 2014). Inactivation of the vBNST by muscimol or norepinephrine antagonists attenuated TMT-induced fear response (Fendt et al., 2003; Fendt et al., 2005). Induction of Fos mRNA in the dlBNST in response to TMT has been previously reported (Asok et al., 2013). In our studies, application of CART peptide in the CeA in the absence of TMT, increased Fos expression in the CeA and vBNST but not in the dlBNST. While multiple sub-regions of the BNST may be involved in fear processing, our study focused on the vBNST as the CeA-vBNST axis is subject to CART-mediated modulation.

The CART peptide is abundantly expressed in the neurons of the CeA while the fiber terminals are seen in the vBNST (Sharma et al., 2014; Upadhya et al., 2013). The capsular CeA and lateral CeA contains CART-positive soma while the medial CeA is devoid of CART-containing cells. However, all three subdivisions of the CeA are rich in CART-positive fibres (Supplementary Fig S2). In the above studies, immunoneutralisation of the CART activity in the CeA was used to demonstrate a functional role for CART in processing innate fear. Further, Fos induction was seen in the CeA as well as vBNST in response to TMT (Sharma et al., 2014). In the present study, immunoneutralization of endogenous CART at the CeA not only reduced the freezing response to TMT but also attenuated the vBNST activation. To underscore the causality of CART activity in inducing freezing, within the framework of the CeA-vBNST, we show that exogenously administered CART intensifies the behavioral response to TMT and concomitantly augments vBNST activation. Further, silencing of the CeA neurons by lidocaine not only attenuated the freezing response to TMT, but also reduced Fos expression in the vBNST. Collectively, the data suggest that the CeA-vBNST axis is central to TMT-induced fear processing. CART signaling modulates the activity of the CeA neurons, which in turn, regulates vBNST activation and fear expression. In principle, unilateral manipulation of the CeA could be used to further dissect the CeA-vBNST circuit. However, the CeA may project to both the ipsilateral and contralateral vBNSTs (Oler et al., 2017) and may confound the analysis.

The detailed connectivity of the CART-modulated CeA-vBNST module is currently unclear. Given that most CeA-vBNST projections are GABA-ergic (Crestani et al., 2013; Dong et al., 2001), the occurrence of monosynaptic connectivity appears unlikely to explain the one to one activity correspondence observed in our studies between CeA and vBNST. Disinhibition via an intermediate interneuron is an alternative, though this is yet to be experimentally determined. A major limitation in mapping the CART-responsive circuitry is the elusive identity of the CART receptor/s.

The CeA-vBNST circuit identified in this study may be an integral component of information flow from the CoA/MeA to the hypothalamus and PAG. It has been shown that the BNST sends afferents to the PAG passing through the anterior hypothalamic nucleus and ventromedial hypothalamus (Dong and Swanson, 2004, 2006). The CeA may have other parallel outputs including direct afferents to the PAG and the laterodorsal tegmental area, both of which are involved in TMT-induced fear (Kessler et al., 2012; Vianna and Brandao, 2003; Yang et al., 2016). This study, together with the previous implication of BNST in TMT-induced fear responses (Fendt et al., 2003; Fendt et al., 2005) emphasizes the CeA-vBNST route as one of the important outputs of the CeA.

In order to understand the CeA-BNST CART circuitry in greater mechanistic detail, high resolution mapping of CART inputs to the CeA and of the projections of the CART-responsive CeA neurons to the sub-regions of the BNST needs to be undertaken in the future. Similarly, specific manipulation of the activity of CART-responsive CeA neurons will circumvent the relatively coarse (spatially) stereotaxy-mediated drug/immunoneutralization approaches. Both these endeavours will benefit immensely from cell type specific transgenic driver lines that will allow regulated expression of tracer molecules or chemo/optogenetic reagents. The choice of the transgenic lines will depend upon the identification of the specific cell types in the CeA that express and/or respond to CART.

### 4.2. CART signaling in the CeA

Our data indicate that CART-signaling modulates the activity of CeA neurons, which, via the vBNST, regulates expression of innate fear. To investigate the mechanism of CART activity in the CeA we investigated the role of NMDA-R-mediated glutamatergic signaling in the CeA neurons. Our results suggest that CART-mediated intensification of the fear response to TMT and augmented activation of the vBNST, are mediated through NMDA-R activity. Consistent with our observation, in mice, a significant proportion of CeA neurons projecting to the vBNST express the NMDA-R in their somato-dendritic compartments (Beckerman and Glass, 2012). Morphine-induced induction of cFos in vBNST neurons is blocked when NMDA-R is genetically silenced in the CeA, implicating functional connectivity between the NMDA-R expressing CeA neurons and the vBNST (Beckerman and Glass, 2012).

It remains unclear whether CART function at the CeA is pre- or postsynaptic. Again, the identification of the CART receptor/s remains a major hurdle in answering this question. CART has been previously found to potentiate NMDA-R activity by promoting phosphorylation of the NR1 subunit in sensory neurons suggesting a postsynaptic function (Chiu et al., 2010). In our studies, injection of CART peptide in the CeA was sufficient to increase Fos induction in both the CeA and vBNST suggesting an enhancement of the baseline excitatory glutamatergic drive in the CeA.

## 5. Conclusion

The CeA-vBNST axis is emerging as a major neural substrate encoding negative valence in physiological and behavioral responses to stress (Avery et al., 2016; Fox et al., 2015; Shackman and Fox, 2016). Our study identifies CART signaling as a major modulator of innate fear gating the information flow in the CeA-vBNST circuit. These studies define a novel mechanistic framework in our understanding of survival instincts subserved by hard-wired circuitry subject to peptidergic modulation.

## ACKNOWLEDGEMENTS

We thank Ms. Geetanjali Nerurkar for maintaining the SD rat colonies. We acknowledge the IISER Pune Microscopy Facility and the National Facility for Gene Function in Health and Disease (NFGFHD) at IISER Pune for providing access to equipment and infrastructure.

